# Angiotensin II Type 2 Receptor Potentiates Skeletal Muscle Satellite Cell Differentiation via the GSK3β/β-catenin Pathway

**DOI:** 10.1101/2022.09.30.510328

**Authors:** C Seth Lott, Bysani Chandrasekar, Patrice Delafontaine, Tadashi Yoshida

## Abstract

Patients with advanced congestive heart failure (CHF) or chronic kidney disease (CKD) often have increased systemic angiotensin II (Ang II) levels and cachexia. We previously demonstrated that Ang II infusion in rodents results in skeletal muscle wasting and reduced muscle regenerative potential via Ang II type 1 receptor (AT1R) signaling, potentially contributing to cachexia in CHF and CKD. Contrary to AT1R signaling, we found that signaling via Ang II type 2 receptor (AT2R) potentiates skeletal muscle satellite cell (SC) differentiation and muscle regenerative potential. However, mechanisms whereby AT2R regulates SC differentiation and cachexia development remain unknown. In this study, we found that GSK3β activity was significantly suppressed during SC differentiation, whereas it was retained in SCs with AT2R knockdown. AT2R knockdown leads to higher GSK3β and decreased β-catenin activities both *in vitro* and *in vivo*. Treatment with GSK3β inhibitor BIO restored β-catenin activity and differentiation capacity of SCs with AT2R knockdown. Conversely, transgenic overexpression of AT2R in SCs inhibited GSK3β, associated with increased β-catenin activity and SC myogenic capacity both *in vitro* and *in vivo*. Interestingly, AT2R expression in undifferentiated SCs was regulated post-transcriptionally. An increase in systemic Ang II blunted AT2R induction during muscle regeneration. However, overexpression of AT2R restored AT2R levels and myogenesis *in vivo*. Together, these data suggest that the AT2R/GSK3β/β- catenin signaling pathway could serve as a potential therapeutic target to promote muscle regenerative capacity in chronic disease conditions characterized by heightened activation of the renin-angiotensin system, such as CHF and CKD.

## Introduction

The renin-angiotensin system (RAS) is a critical regulator of blood volume and systemic vascular resistance. Classically, the RAS is considered as a circulating hormonal system, with angiotensin II (Ang II) acting as a main bioactive peptide. However, it has become clear that a local “tissue RAS” is present in most organs and functions as an endocrine, paracrine and intracrine system^1,2^.

Studies from our group and others have shown that Ang II plays an important role in maintaining skeletal muscle homeostasis. Ang II infusion in rodents caused skeletal muscle atrophy through multiple mechanisms, including activation of the ubiquitin–proteasome system (UPS)^3,4^, reduction of insulin-like growth factor-1 (IGF-1)^3,5,6^, disruption of energy balance (decreased ATP levels)^7,8^, decreased food intake (limited availability of nutrients)^9^, and increased mitochondrial damage^10^. These atrophying effects of Ang II may be relevant to development of cachexia (muscle wasting) in patients with chronic diseases. Patients with advanced congestive heart failure (CHF) and chronic kidney disease (CKD) often have elevated circulating levels of Ang II, and treatment with an angiotensin converting enzyme inhibitor (ACEi) can reduce weight loss^11^. Chronic increase in systemic Ang II has also been shown to play a role in the development of cachexia during cancer^12^ and chronic obstructive pulmonary disease (COPD)^13^. Penafuerte *et al*.^14^ reported that plasma Ang II levels are significantly increased in pre-cachectic and cachectic cancer patients and proposed Ang II as a blood biomarker and the master upstream regulator of pre-cachexia.

Muscle damage, specifically to the muscle membrane (sarcolemma) is a characteristic feature of muscle atrophy. Although muscle damage normally activates skeletal muscle satellite (stem) cells (SCs) to trigger muscle regeneration, many of the chronic disease conditions, such as disuse^15–18^, denervation^19^, COPD^20–25^, CKD^26,27^, burn injury^28,29^, diabetes^30–35^ and cancer^36–40^, are associated with reduced regeneration.

Similar to these chronic disease conditions, our studies have provided evidence that the RAS suppresses skeletal muscle regeneration, potentially contributing to development of muscle atrophy. We have found that the two types of Ang II receptors, type 1 (AT1R) and type 2 (AT2R), regulate different stages of SC differentiation. AT1R is predominantly expressed in quiescent and proliferating SCs^41^, whereas AT2R expression is strongly induced during differentiation into myotubes^42,43^. AT1R activation inhibits SC proliferation, leading to depletion of the SC pool and reduced muscle regenerative capacity. In contrast, AT2R activation likely potentiates SC differentiation; AT2R knockdown inhibits SC differentiation and skeletal muscle regeneration. Importantly, it has been shown that AT2R expression is robustly increased in many tissues/cells during embryogenesis, but falls below detectable levels in many adult tissues^44–47^, suggesting that AT2R may regulate differentiation of multiple cell types in addition to SCs. Therefore, targeting the AT2R signaling pathway has the potential to lead to the development of novel interventions to restore lowered skeletal muscle regenerative capacity in chronic diseases and aging, and to provide insights into the molecular mechanisms that regulate differentiation of multiple cell lineages. However, the downstream signaling pathways of AT2R in SCs, and the molecular mechanisms whereby AT2R regulates SC differentiation are not known. Here, we aimed to identify novel AT2R downstream signaling pathways that regulate SC differentiation processes.

## Materials and Methods

### Animals

Animal protocols were approved by Animal Care and Use Committee at Tulane University and the University of Missouri-Columbia. 8- to 10-week-old male C57BL/6 mice (The Jackson Laboratory) were used in this study. Cardiotoxin (CTX, 10 μM) was injected at 4 different locations in tibialis anterior muscles (a total of 25 μl) or 6 different locations in gastrocnemius muscles (50 μl). In order to overexpress AT2R in cell type-specific manner, we generated CAG-CAT^flox^-AT2R mice. These transgenic mice have a reporter gene chloramphenicol acetyltransferase (CAT) flanked by two loxP sites (CAT^flox^) connected under the CAG promoter (CMV early enhancer element, chicken β-actin gene promoter and the splice acceptor of the rabbit β-globin gene), followed by mouse AT2R coding region. Due to the presence of a stop codon under the CAT reporter, AT2R expression is not induced from CAG-CAT^flox^-AT2R allele. To generate CAG-CAT^flox^-AT2R mice, AT2R ORF sequence was amplified with primers GCC GGA TCC GCC GCC ACC ATG AAG GAC AAC TTC AGT TTT GCT GC and GCC GGA TCC TTA AGA CAC AAA GGT GTC CAT TTC TC, which add Kozak sequence^48^ (GCC GCC ACC) in front of the start codon and BamHI recognition sites at both ends. The amplified sequence was subcloned into CAG-lox-CAT-lox vector(gift from Jeffrey Robbins; Addgene plasmid # 53959)^49^ using BamHI cloning site. The plasmid was digested with SalI, NotI and PvuI, and ~4.8 kbp DNA fragment was purified. The DNA fragment was injected into C57BL/6 mouse oocytes at University of Missouri Animal Modeling Core, and four founder lines (F888, F898, F900 and F902) were obtained. Pax7^CreER^ mice (B6.Cg-Pax7^tm1(cre/ERT2)Gaka^/J; The Jackson Laboratory #017763)^50^ and R26^LSL-EYFP^ mice(B6.129X1-Gt(ROSA)26Sor^tm1(EYFP)Cos^/J; The Jackson Laboratory #006148)^51^ were obtained from Jackson Laboratory. Southern blot probe for CAG-CAT^flox^-AT2R mice was generated using PCR DIG probe synthesis kit (Roche) with primers GCC GAA TTC GCC GCC ACC ATG AAG GAC AAC TTC AGT TTT GCT GCC and GCA GGT AAT AAA AAA ATA TGC TTG CAA ACA TGT TCA G. The probe is located within exon 3 of the AT2R. For routine PCR genotyping, tail DNA was analyzed by primers AGG CCC TAC AGG TTG TCT TCC CAA CTT GCC CCT TGC TCC and CCA CCC GAG AAA TAT GCT CAG TGG TCT GCT GGG ATT GCC, with additional internal control primers CTA GGC CAC AGA ATT GAA AGA TCT (oIMR7338; The Jackson Laboratory) and GTA GGT GGA AAT TCT AGC ATC ATC C (oIMR7339; Jackson Laboratory). Ang II (Phoenix Pharmaceuticals) was infused via subcutaneously implanted osmotic minipump (ALZET model 1007D) to continuously deliver 1.5 μg/kg/min, as described previously^4^.

### Antibodies and reagents

The following antibodies were used in immunoblotting and immunohistochemistry: anti-AT2R and anti-laminin were from Millipore-Sigma, anti-myogenin was from Abcam, and anti-desmin, anti-GSK3β^Ser9^/GSK3β, anti-Akt^Ser473^/Akt were from Cell Signaling. Tamoxifen (Sigma-Aldrich C8267) was dissolved in corn oil (20 mg/ml) and injected IP at the dose of 75 mg/kg once daily for 5 days, and experiments were performed 1 week after the last injection. BIO ((2’Z,3’E)-6-Bromoindirubin-3’-oxime; GSK3β inhibitor) and methyl-BIO (MBIO, control) were obtained from Millipore, and a 10 mM stock solution was prepared in DMSO. Cells were treated with BIO/MBIO at 2 μM concentration. A stock solution of recombinant mouse Wnt3A (R&D systems) was prepared at 40 μg/ml in 0.1% BSA/PBS and used between 10-100 ng/ml concentration in culture.

### Cell culture

SCs were collected as previously described ^52,53^. Briefly, hindlimb muscles (gastrocnemius, soleus, tibialis anterior and extensor digitorum longus) were incubated with 0.2% (w/v) collagenase type II in DMEM and 10% horse serum for 2 h at 37°C. Digested muscles were dissociated by triturating with Pasteur pipette, washed and further digested with 1 U/ml dispase for 1 h at 37°C. Cells were filtered, washed and incubated in proliferation medium (DMEM supplemented with 20% FBS (Mediatech); 10% HS; 2% chicken embryo extract (US Biological); 1 mM sodium pyruvate; 2 mM L-glutamine and penicillin/streptomycin) for 40 min to let the fibroblasts attach onto the plate and unattached cells were plated onto Matrigel-coated plates in the proliferation medium at the density of 1×10^4^ cells/cm^2^. C2C12 myoblasts (ATCC) were cultured in C2C12 growth medium (DMEM supplemented with 20% FBS, 1 mM sodium pyruvate and 2 mM L-glutamine). After SCs reached confluence, cells were switched to the differentiation medium (DMEM supplemented with 2% horse serum, 1 mM sodium pyruvate and 2 mM L-glutamine) to induce differentiation. AT2R siRNA (ThermoFisher s201015) and negative control (ThermoFisher 4390844) were transfected into cells using Lipofectamine RNAiMAX (ThermoFisher) following manufacturer’s instruction.

### Quantitative RT-PCR

Quantitative RT-PCR (qRT-PCR) was performed using TaqMan assays except for AT2R, and probes used in this study were obtained from ThermoFisher. Hprt was used as an internal control. Pax7: Mm01354484_m1; MyoD: Mm00440387_m1; Myogenin: Mm00446194_m1; Myh3: Mm01332463_m1; Mymk: Mm00481256_m1; Mymx: Mm04932812_g1; Follistatin: Mm00514982_m1; Hprt: Mm03024075_m1. AT2R qRT-PCR was performed using RT^2^ qPCR Primer Assay (Qiagen). AT2R: PPM04811A; Hprt: PPM03559F.

### Immunohistochemical staining and analysis of myofiber cross sectional area

Myofiber cross sectional area (CSA) was determined on H/E or laminin-stained paraffin sections (6 μm) as described previously^42^. Images were taken using Cytation 5 Cell Imaging Multimode Reader (Biotek), and analyzed by ImageJ/Fiji software (Ver. 2.3.0/1.53q).

### Phosphoprotein Profiling by the Phospho Explorer Antibody Microarray

The PhosphoExplorer Antibody Array (Cat# PEX100, Full Moon BioSystems) contains 1,318 site-specific and phospho-specific antibodies, each of which has two replicates. This is an ELISA based phosphorylation assay for qualitative protein phosphorylation profiling. The microplate also contains multiple positive and negative controls. Assay was performed by Full Moon BioSystems using the protein lysate prepared from primary cultured SC following their established protocol.

### TCF/LEF reporter assay

M50 Super 8x TopFlash reporter vector^54^ contains 7 TCF/LEF binding sites upstream of firefly luciferase gene, allowing quantification of TCF/LEF activation. pGL4.74[hRluc/TK] vector (Promega) was co-transfected, and Renilla luciferase activity was used to normalize the firefly luciferase signal. *In vitro* luciferase reporter assays were performed in 24 well culture plates. Cells were plated at 1.67×10^5^ cells/well, and plamids (500 ng of M50/M51 and 25 ng of pGL4.74[hRluc/TK] per well) were transfected using Lipofectamine 3000 reagent (ThermoFisher). *In vivo* luciferase reporter assay in hindlimb muscles were performed by electroporating the plasmids (20 μg of M50/M51 and 1 μg of pGL4.74[hRluc/TK] per TA muscle). Firefly and Renilla luciferase activities were measured by Dual-Luciferase Reporter Assay System (Promega) following manufacture’s instructions.

### Plasmid electroporation to hindlimb muscles

Mice were anesthetized with a mixture of 75 mg/kg ketamine 15 mg/kg xylazine hydrochloride administered by IP injection. Mice hindlimbs were shaved, and plasmids (1 μg/μl in saline, 20 μg per muscle) were injected into TA muscle with a 22G needle (Hamilton model 705). Transcutaneous pulses were applied by two stainless steel plate electrodes (Caliper Electrode model 384, BTX) with a 0.5 cm distance between the two plates. The electrical conduct was ensured using a conductive gel. Electric pulses with a standard square wave were delivered by an electroporator (ECM830 Electro Square Porator, BTX). Eight pulses of 50 V/cm were administered to the muscle with a delivery rate of 1 pulse/sec with each pulse 20 msec in duration. Muscles were collected at indicated time points.

### RNA-seq data analysis

RNA-seq dataset GSE53398 was retrieved from the NIH Gene Expression Omnibus (GEO) database. The quality of sequences was analyzed by FastQC before and after trimming and adaptor clipping by Trimmomatic^55^. The trimmed sequences were aligned to mouse genome by HISAT2^56^ using GRCm38 index, followed by sorting alignments by SAM tools^57^ and read counting by featureCounts^58^. The count data were further processed by R (version 4.2.1) with libraries edgeR^59^ and limma^60^.

### Statistical analysis

All data represent mean ± SEM, and results were analyzed by 2-way or 3-way ANOVA depending on the number of independent variables, followed by Tukey’s honest significance (HSD) test for multiple comparison.

## Results

### Phosphoprotein changes downstream of AT2R in satellite cells

It was previously shown that MAPK signaling plays an important role in C2C12 myoblast differentiation^61^ and AT2R signaling attenuates ERK1/2 activity in vascular smooth muscle cells^62^. Our previous data show that during C2C12 myoblast to myotube differentiation, the phosphorylation of ERK1/2 was down-regulated at the initial stage (day 1), followed by an increase above the level of predifferentiation (day 2-3). Further, AT2R knockdown increases ERK1/2 phosphorylation throughout this bi-phasic phase of ERK1/2 regulation, and inhibition of ERK1/2 by a MEK inhibitor restored C2C12 differentiation^42^. These data strongly suggest that AT2R positively regulates myoblast differentiation via down-regulation of ERK1/2 signaling. However, there is no further information regarding AT2R downstream signaling in SCs. To further explore AT2R downstream signaling, we analyzed AT2R-mediated phosphoprotein changes in SCs. We transfected SCs with AT2R-specific siRNA and induced the cells to differentiate for 2 days. As reported previously^42^, we observed efficient AT2R knockdown and suppression of myogenesis in these cells **(Fig. 1A, inset)**. We analyzed phosphoprotein changes using a Phospho-Explorer Antibody Microarray (FullMoon Biosystems), in which 1,318 unique antibodies against phosphoproteins and corresponding total proteins are spotted. Among these proteins, 138 proteins (71 increased and 67 decreased) showed altered phosphorylation levels in AT2R silenced SCs compared to control above the >2 fold threshold **(Fig. 1A)**. These phosphoprotein changes were analyzed by Qiagen Ingenuity Pathway Analysis software (Ver. 76765844), and the top canonical pathways and diseases and functions annotations were obtained **(Fig. 1B, C)**. Among the top canonical pathways were TGF-β and Wnt/β-catenin signaling pathways, both of which play important roles in skeletal muscle regeneration and fibrosis development. TGF-β and Wnt pathways are amongst the most well-studied cell signal transduction pathways in metazoan biology, and TGF-β can antagonize or synergize with Wnt signaling, in a context dependent manner^63,64^. Fig. 1D shows changes in GSK3β and β-catenin phosphorylation, both of which are critical components in the Wnt signaling pathway. Phosphorylation of GSK3β at Ser216 and Ser9 was decreased in AT2R-silenced SCs. In contrast, AT2R knockdown increased β-catenin phospholyration at Thr41/Ser45, Ser37, Ser33, Tyr489, but not at Tyr654. Since dephosphorylation of GSK3β results in the activation of its kinase activity, which in turn will lead to phosphorylation of β-catenin and subsequent degradation, our data suggest that silencing AT2R inhibits the Wnt/GSK3β/β-catenin pathway in SCs.

**Fig. 1.**
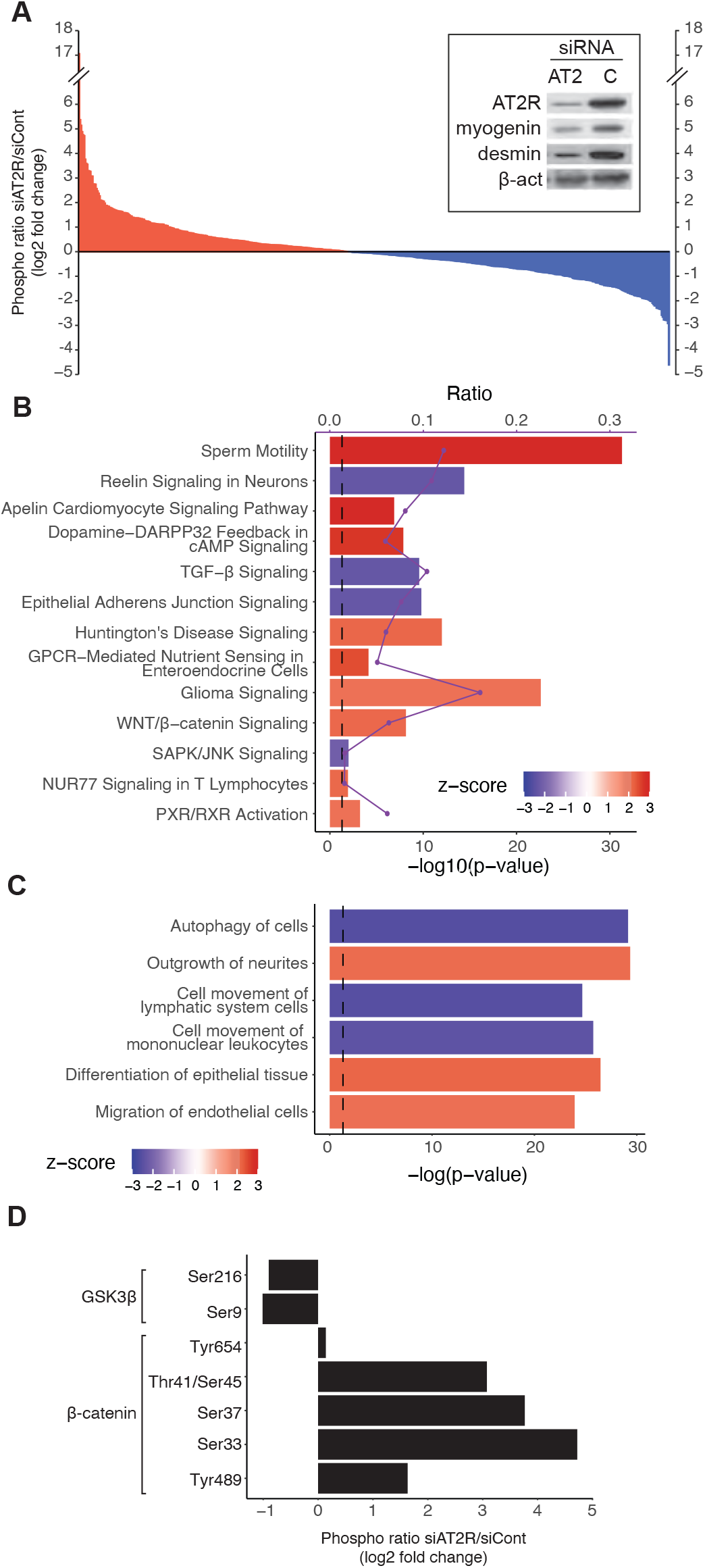
Phosphoprotein profiling in AT2R silenced muscle satellite cells. **(A)** Phosphoprotein changes analyzed by PhosphoExplorer array in muscle satellite cells (SCs) transfected with AT2R or control siRNA, followed by induction of differentiation for 2 days. Log2 fold change values of AT2R siRNA-transfected cells compared to control are shown, with increased phosphorylation in red and decreased phosphorylation in blue. **(B, C)** IPA Canonical Pathway analysis (B) and Diseases and Functions Annotations (C) for the phosphoprotein changes with absolute log2 fold change values over 1 (i.e. >2 fold change). Bars indicate -log_10_p-value, and lines show the ratio of the proteins that were changed among the proteins associated with each pathway. The red and blue colors of the bar chart indicate activation z-score, and only the pathways with absolute z-scores higher than 2 (i.e. considered statistically significant). **(D)** Changes in phosphorylation levels in GSK3β (Ser216 and Ser9), and β-catenin (Tyr654, Thr41/Ser45, Ser37, Ser33 and Tyr489).

### AT2R regulates the GSK3β/β-catenin pathway in satellite cells

The Wnt/GSK3β/β-catenin pathway interacts with multiple other signaling pathways, including the PI3K/Akt pathway^65^. We analyzed changes in Akt and GSK3β phosphorylation in SCs following AT2R knockdown **(Fig. 2A, B)**. Consistent with published reports^66–68^, our results show that phosphorylation of Akt at Ser473 and GSK3β at Ser9 was increased during SC differentiation. Since phosphorylation at Ser9 is considered inhibitory, these data indicate inhibition of GSK3β kinase activity. Notably, silencing AT2R significantly blocked p-Akt^Ser473^ and p-GSK3β^Ser9^, indicating that GSK3β is activated. To further analyze GSK3β activation, T cell factor/lymphoid enhancer factor family (TCF/LEF) reporter plasmid (Super TOPFlash)^54^ was transfected into primary SCs to measure changes in GSK3β/β-catenin signaling in AT2R silenced cells **(Fig. 2A)**. Consistent with the increased Akt^Ser473^ and GSK3β^Ser9^ phosphorylation, the TCF/LEF-dependent reporter activity was increased during SC differentiation, and this effect was reversed by AT2R knockdown, suggesting that AT2R-mediated inhibition of Akt/GSK3β signaling leads to potentiation of β-catenin activity.

**Fig. 2.**
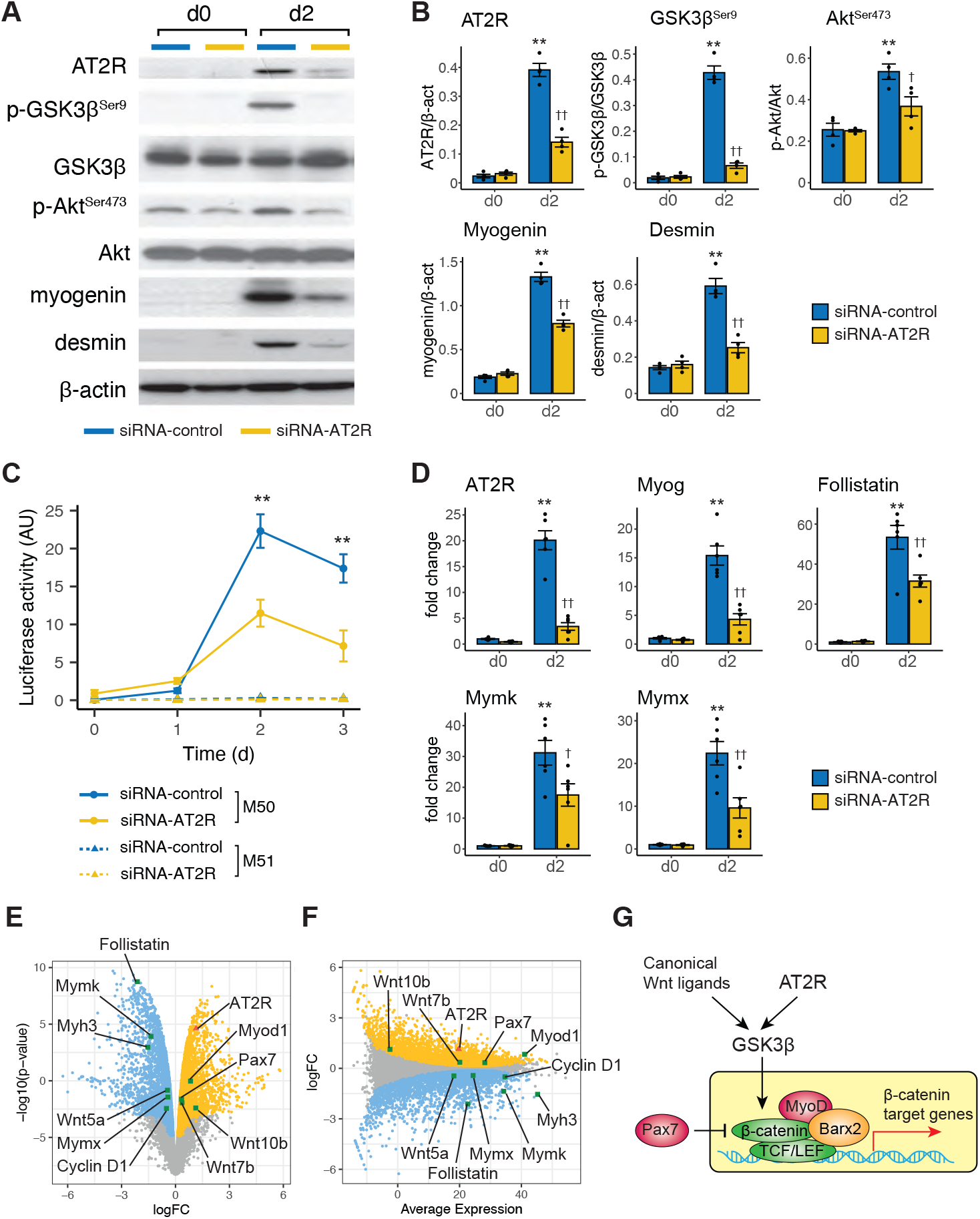
AT2R knockdown in SCs inhibits Akt/GSK3β/β-catenin pathway and myogenic differentiation. **(A)** Immunoblot analysis of Akt/GSK3β pathway and myogenic differentiation markers. Primary SCs were transfected with AT2R or control siRNA, induced to differentiate for 2 days, and protein expressions were analyzed by immunoblotting. **(B)** Quantification of the immunoblots from (A). N=4, mean±SE, *p<0.05, **p<0.01. **(C)** Quantification of TCF/LEF reporter activity. Primary cultured SCs were transfected with TOPFlash reporter vector M50 or mutant control vector M51, together with control or AT2R siRNA. Cells were induced to differentiate, and luciferase activity was measured at the indicated time points. Firefly luciferase activity from M50/M51 vectors was normalized by *Renilla* luciferase activity from the control pGL4.74[hRluc/TK] vector. **(D)** From the same set of samples as in (A), mRNA expression of myogenic differentiation markers and β-catenin target genes was analyzed by qRT-PCR. **(E, F)** Analyses of RNA-seq dataset of primary myoblasts from WT and Barx2-KO mice (GEO accession: GSE53398). mRNA expression of each gene is visualized in volcano plot (E) and Mean-Difference (MD) plot (F). Genes in yellow and blue indicate increased and decreased expression, respectively, in Barx2 deleted myoblasts compared to WT control. Expression of SC markers (Pax7, MyoD, Myog, Myh3), Wnt ligands (canonical: Wnt7b and Wnt10a; non-canonical: Wnt5a), Wnt target genes (Follistatin, Cyclin D1, Mymk, Mymx) and AT2R are shown in the plots. **(G)** Hypothetical model of Wnt/GSK3β/β-catenin and AT2R signaling pathways. Barx2, Pax7, MyoD and β-catenin form Wnt effector complex and regulate TCF/LEF-mediated transcriptional activation, including myogenic genes such as Mymk and Mymx. AT2R activates GSK3β downstream of, or independently of Wnt ligands. In (B) and (D), asterisks (*) and daggers (†) indicate statistical significance between control siRNA (d0) vs. control siRNA (d2) and control siRNA (d2) vs. AT2R siRNA (d2), respectively. In (C), asterisks (*) indicate statistical significance between control siRNA vs. AT2R siRNA at d2 or d3. N=4 in (B) and (C), and N=6 in (D). Error bars show SEM in all the figures. *p<0.05; **p<0.01; †p<0.05; ††p<0.01.

Jones *et al*.^69^ have previously demonstrated that canonical Wnt/β-catenin signaling primes myoblasts to differentiate by stimulating myogenin and follistatin expression. Mechanistically, myogenin binds follistatin promoter to activate its transcription. In AT2R silenced SCs, we found that both myogenin and follistatin mRNA levels were decreased, further suggesting that AT2R activates SC differentiation via β-catenin signaling **(Fig. 2D)**.

Zhuang *et al*.^70^ found that Barx2 homeobox protein, Pax7, and MyoD form a Wnt effector complex, and play a role in SC differentiation. Canonical Wnt signaling induces Barx2 expression in SCs and perturbation of Barx2 leads to misregulation of Wnt target genes. It is suggested that Barx2 forms a transcriptional activator complex with β-catenin, TCF/LEF factors and MyoD, promoting transcription via TCF/LEF sites. Pax7 inhibits the function of this complex, likely via interaction with β-catenin. To gain further insights into the potential involvement of AT2R in canonical Wnt/GSK3β/β-catenin signaling in SCs, we analyzed RNA-seq dataset from Barx2 knockout SCs (GSE53398^70^, **Fig. 2E, F**). In Barx2 knockout SCs, β-catenin target genes such as Mymk, Mymx, Follistatin and Cyclin D1 were downregulated. Expression of Pax7 and MyoD, which form a complex with Barx2, and canonical Wnt ligands Wnt7b and Wnt10b were increased, whereas non-canonical Wnt5a was decreased. Pax7, MyoD, Wnt7b and Wnt10b act upstream or together with Barx2, and the increased expression of these genes is likely due to a feedback mechanism. Interestingly, AT2R expression was also increased in Barx2 silenced SCs, suggesting that AT2R acts upstream of Barx2/β-catenin complex **(Fig. 2G)**.

### Inhibition of GSK3β restores myogenic capacity of AT2R silenced SCs

To gain further insight into how AT2R regulates the Akt/GSK3β/β-catenin pathway, we aimed to restore AT2R downstream signaling in AT2R silenced SCs by (1) constitutively-active Akt (caAkt), (2) GSK3β inhibitor, and (3) Wnt ligand. Co-transfection of caAkt (myrAktΔ4-129) and AT2R siRNA increased phosphorylation of GSK3β at Ser9 (i.e. GSK3β was inhibited) and restored SC myogenic potential **(Fig. S1A, B)**. Moreover, consistent with our hypothesis, TCF/LEF reporter activity was also restored in these cells **(Fig. S1C)**. Treatment of the cells with GSK3β inhibitor BIO ((2’Z,3’E)-6-Bromoindirubin-3’-oxime) increased GSK3β^Ser9^ phosphorylation (i.e. inhibition of GSK3β) in AT2R silenced cells **(Fig. 3A, B)**. Importantly, BIO treatment restored SC differentiation, indicated by increased myogenin and desmin expression. Consistent with the increased GSK3β^Ser9^ phosphorylation, BIO restored TCF/LEF reporter activity **(Fig. 3E)**, and myofiber generation **(Fig. 3G)** in AT2R silenced cells. In contrast to caAkt and BIO, Wnt3A did not alter GSK3β^Ser9^ phosphorylation **(Fig. 3C, D)**. Indeed, myogenin and desmin expression indicated that Wnt3A Consistently failed to restore SC myogenic capacity **(Fig. 3C, D)**, and myofiber generation **(Fig. 3H)**. Furthermore, TCF/LEF reporter assay showed that Wnt3A did not restore β-catenin activity **(Fig. 3F)**. These data suggest that AT2R regulates the GSK3β/β-catenin pathway downstream of, or independent of the Wnt pathway.

**Fig. 3.**
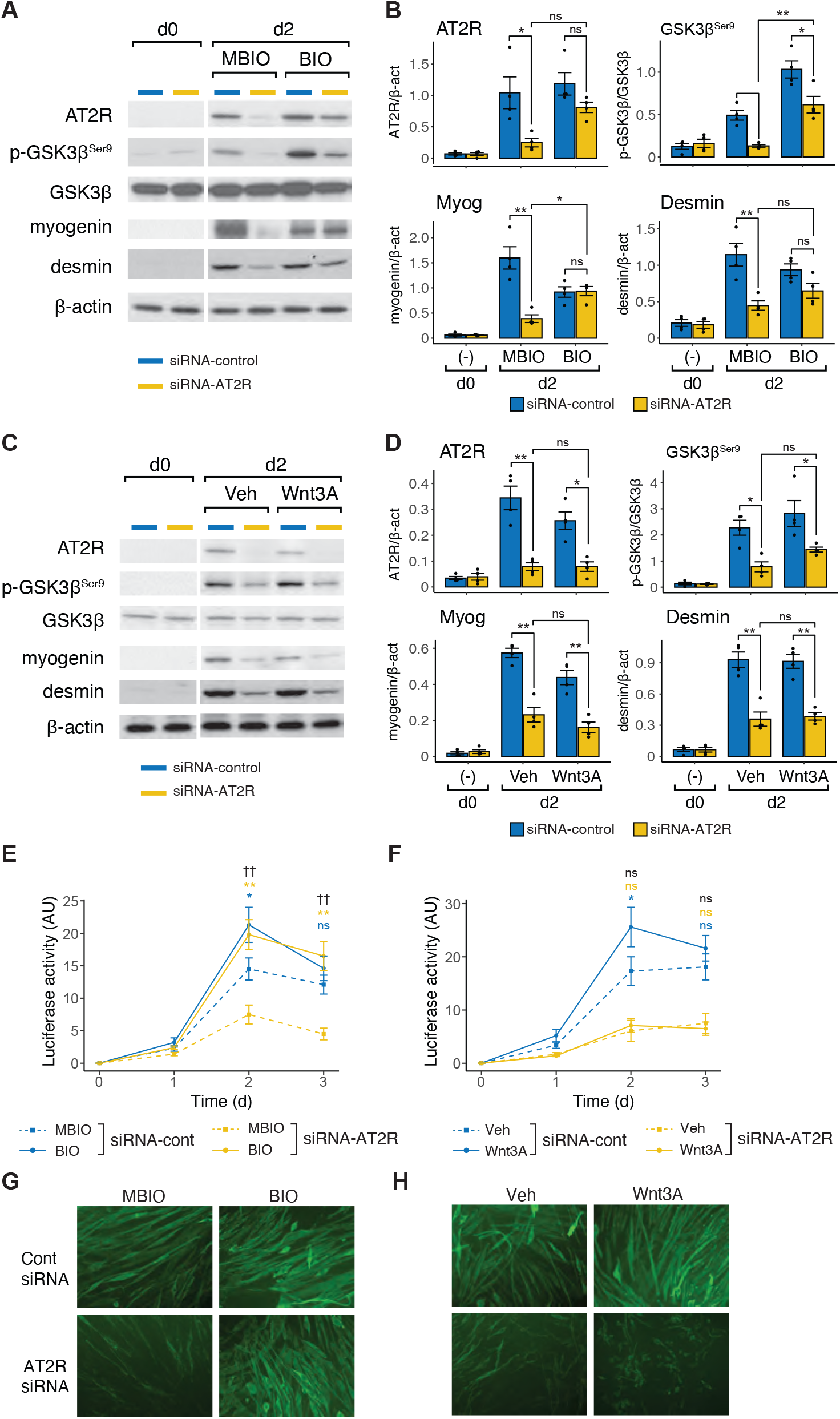
Restoration of AT2R knockdown-mediated SC differentiation by GSK3β inhibition, but not by Wnt ligand. **(A-D)** Immunoblot analysis of GSK3β and myogenic differentiation markers. Primary SCs were transfected with AT2R or control siRNA, induced to differentiate for 2 days with or without the presence of GSK3β inhibitor BIO (2 μM; A, B) or Wnt3A (30 ng/ml; C, D). Quantification of the Immunoblot data in (A) and (C) are shown in (B) and (D), respectively. **(E, F)** Quantification of TCF/LEF reporter activity. Primary cultured SCs were transfected with TOPFlash reporter vector M50 or mutant control vector M51, together with AT2R or control siRNA. Cells were induced to differentiate with or without the presence of GSK3β inhibitor BIO (2 μM; E) or Wnt3A (30 ng/ml; F), and luciferase activity was measured at the indicated time points. Firefly luciferase activity from M50/M51 vectors was normalized by *Renilla* luciferase activity from the control pGL4.74[hRluc/TK] vector. **(G, H)** From the same set of samples as in (A) and (C), myofiber development was analyzed by MYH3 staining 2 days after induction of differentiation. (B, C) N=4; mean±SEM; *p<0.05; **p<0.01. (E, F) Asterisks (*) indicate statistical significance between treatments (MBIO/BIO or Veh/Wnt3A) within the same siRNA group (control siRNA: blue; AT2R siRNA: yellow). Daggers (†) indicate statistical significance between treatments (MBIO/BIO or Veh/Wnt3A) in AT2R siRNA treated cells. N=4; mean±SEM; *p<0.05; **p<0.01; †p<0.05.

### Generation of AT2R Tg mice

To investigate whether AT2R activation potentiates SC differentiation *in vivo*, we generated CAG-CAT^flox^-AT2R transgenic mice which overexpress the AT2R gene in selected cell types **(Fig. 4, S2A)** after crossing with select Cre mice. Four founder lines were obtained with different copy numbers of the transgene **(Fig. S2B)**, and the line F902 was selected based on the expression of AT2R transgene (data not shown), and crossed with Pax7^CreER^ mice^50^ and R26^LSL-EYFP^ mice^51^. In Pax7^CreER/+^ mice, CreER (tamoxifen-inducible cre recombinase) is expressed under the control of the Pax7 promoter. In R26^LSL-EYFP^ mice, the EYFP gene is knocked into the ROSA26 (R26) locus with upstream floxed stop codon (LSL), and Cre-mediated recombination allows EYFP to be expressed in specific cell types. Therefore, in this triple-transgenic strain (named SC-CAG-AT2R), AT2R expression is induced in SCs upon tamoxifen administration, and AT2R-overexpressing SCs and their progenies are visualized by EYFP fluorescence **(Fig. 4A and Fig. S2C)**. As a control, SC-EYFP mice (Pax7^CreER/+^; R26^LSL-EYFP^) were used. After tamoxifen administration, SCs and other tissues were collected from SC-CAG-AT2R mice and analyzed for AT2R mRNA expression by qRT-PCR **(Fig. 4B)**. Induction of AT2R mRNA was observed specifically in SCs in these mice, indicating the successful generation of the desired animal strain. Importantly, body weight and myofiber cross sectional area were not altered in SC-CAG-AT2R mice compared to control between 8 and 24 weeks of age **(Fig. S3A-C)**, suggesting that AT2R mRNA overexpression in SCs *in vivo* did not activate a muscle regeneration program under basal conditions.

**Fig. 4.**
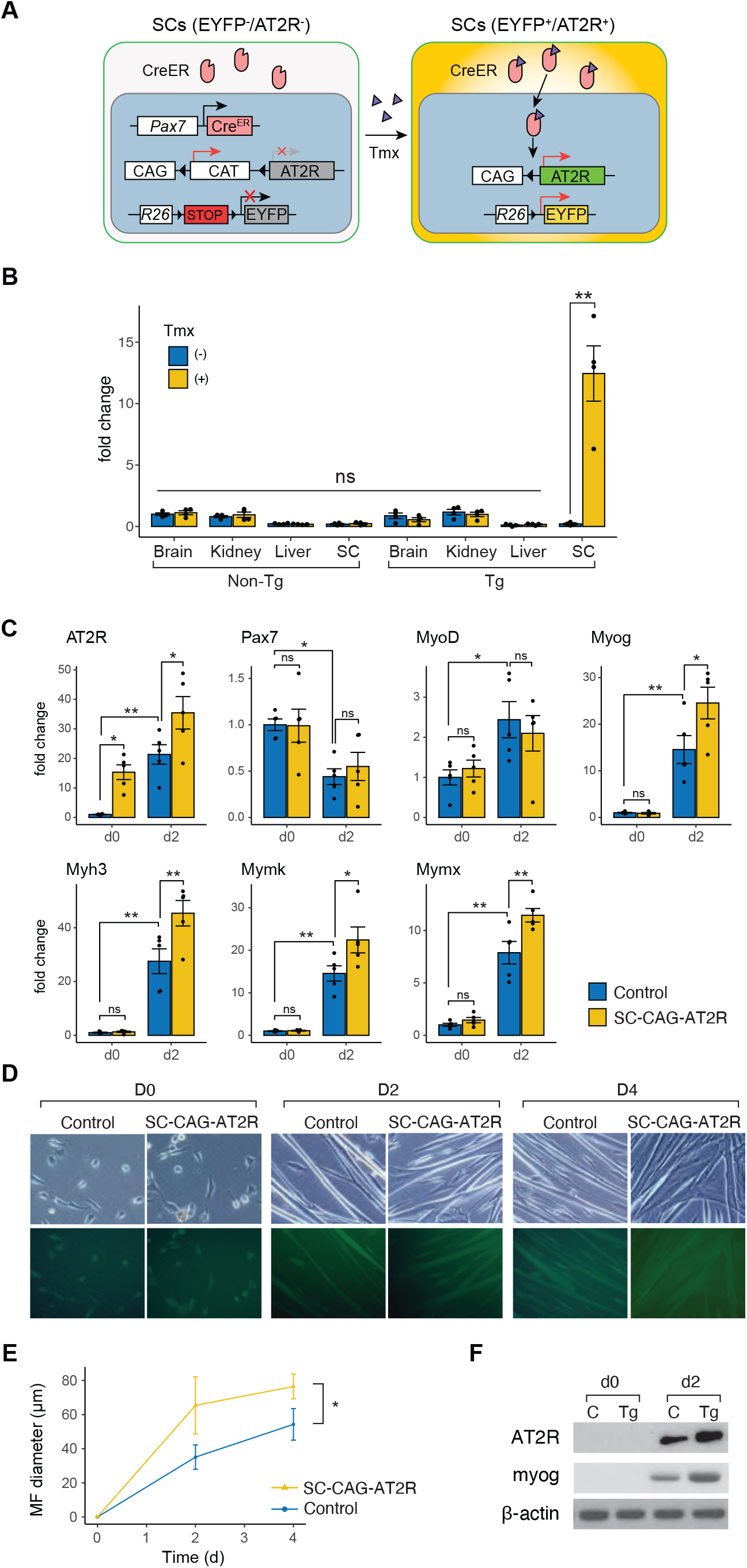
Transgenic AT2R overexpression in SCs potentiates SC differentiation. **(A)** SC-CAG-AT2R animal model. Left panel: In SC-CAG-AT2R (Pax7^CreER/+^; CAG-CAT^flox^-AT2R; R26^LSL-EYFP^) mice, neither AT2R nor EYFP is expressed due to the presence of stop codons upstream of AT2R or EYFP. CreER is expressed specifically in SCs under the control of Pax7 promoter but inactive in the absence of tamoxifen (Tmx). Right panel: Upon tamoxifen administration, CreER becomes active and stop codons are removed via Cre-mediated recombination, allowing AT2R or EYFP to be expressed specifically in SCs. **(B)** Various tissues and SCs were harvested from control (Pax7CreER/+; R26^LSL-EYFP^) and SC-CAG-AT2R mice ±Tmx administration. AT2R mRNA expression was quantified by qRT-PCR. N=4; mean±SEM; *p<0.05. **(C)** Primary SCs were collected from SC-CAG-AT2R and control SC-EYFP mice, cultured *in vitro*, and induced to differentiate for 2 days. Cells were harvested before (d0) and after 2 days of differentiation, and gene expression was analyzed by qRT-PCR. N=5; mean±SEM; *p<0.05; **p<0.01. **(D, E)** SCs were collected as in (C), and myofiber formation was visualized in bright field (top panels in D) and by EYFP fluorescence (bottom panels in D), and myofiber diameter was quantified (E). N=4; mean±SEM; *p<0.05. (F) SCs were collected as in (A), and AT2R and myogenin protein expression was measured by immunoblotting.

### AT2R overexpression increases SC myogenic capacity

SCs were collected from SC-CAG-AT2R mice after tamoxifen administration, and induced to differentiate in culture **(Fig. 4C)**. As expected, AT2R mRNA was higher in SC-CAG-AT2R-derived SCs compared to control before induction of differentiation (d0). At this time point, an increase in AT2R mRNA had no effect on SC marker expression; there was no significant difference in Pax7, MyoD, myogenin, Myh3, Mymk and Mymx expression. However, endogenous AT2R expression was increased upon induction of SC differentiation, and SC-CAG-AT2R transgene had an additive effect on AT2R expression. In contrast to the d0 time point, expression levels of myogenin, Myh3, Mymk and Mymx were markedly increased in SC-CAG-AT2R-derived SCs compared to control at d2, while Pax7 and MyoD expression was not significantly altered **(Fig. 4C)**. Furthermore, an increase in myofiber size was observed upon induction of differentiation in SC-CAG-AT2R-derived SC culture compared to control **(Fig. 4D, E)**. These results suggest that an increase in AT2R expression potentiates SC differentiation into myotubes. Surprisingly, though AT2R mRNA levels were increased ~20 fold, its protein levels were not altered in SC-CAG-AT2R-derived SCs at d0 **(Fig. 4F)**. We did, however, observe an increase in AT2R protein expression in SC-CAG-AT2R-derived SCs following differentiation. These data suggest post-transcriptional suppression of AT2R protein expression (e.g. suppression of translation or enhanced protein degradation) in non-differentiating SCs, and SC differentiation creates an environment in which AT2R protein can be expressed to potentiate SC differentiation. Although it is unknown how AT2R protein levels were suppressed in non-differentiating SCs, these data strongly suggest a mechanism by which AT2R protein levels are strictly controlled in SCs to block precocious activation of SC differentiation.

### Increased skeletal muscle regeneration in SC-CAG-AT2R mice

We next investigated if overexpression of AT2R in SCs in SC-CAG-AT2R mice alters their differentiation and skeletal muscle regeneration *in vivo*. After CTX injury, the size, but not the number, of regenerating myofibers increased in SC-CAG-AT2R mice **(Fig. 5A-C)**. During regeneration, AT2R mRNA expression was increased as reported earlier^42^, with greater AT2R expression in SC-CAG-AT2R mice at all stages **(Fig. 5D)**. Moreover, the increase in AT2R mRNA levels were associated with an increase in the expression levels of Myog, Mymk, Mymx and Myh3 in injured muscles, whereas Pax7 and MyoD expression remained unaffected. It is of note that an increase in AT2R expression at d0 (without CTX injury) did not induce changes in expression of any of the SC markers, suggesting that an increase in AT2R mRNA expression itself does not alter the SC differentiation program.

**Fig. 5.**
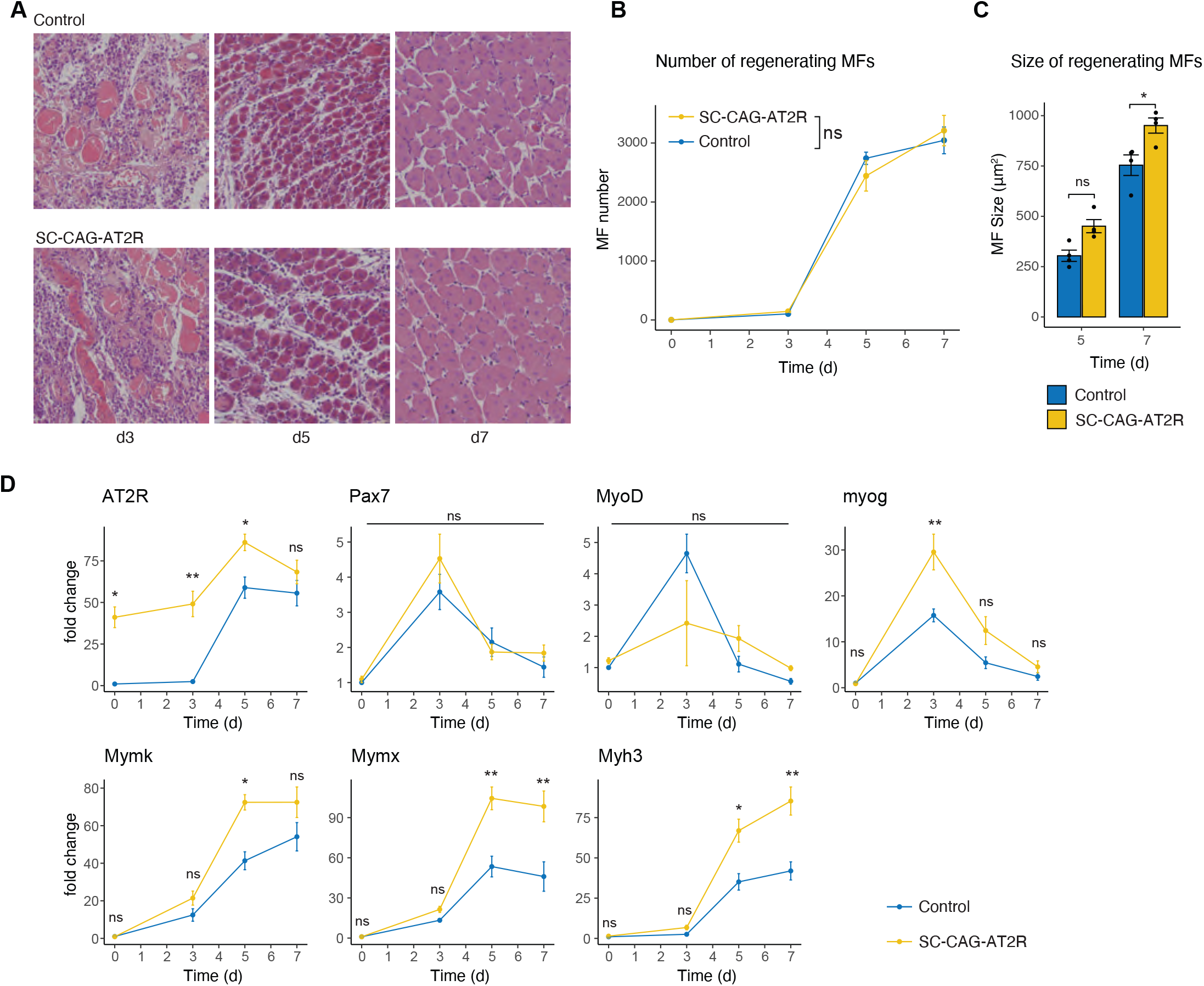
AT2R overexpression in SCs potentiates skeletal muscle regeneration. **(A-C)** H/E images (A) of gastrocnemius muscles from SC-CAG-AT2R and control SC-EYFP mice after CTX injury. The number (B) and size (C) of regenerating myofibers were quantified from these muscles. N=4; mean±SEM; *p<0.05. **(D)** CTX was injected into TA muscles of SC-CAG-AT2R and control mice, and gene expression was analyzed by qRT-PCR at indicated time points.

### AT2R downstream signaling in SCs from SC-CAG-AT2R mice

To study AT2R downstream signaling *in vivo*, SCs were harvested from SC-CAG-AT2R and control (SC-EYFP) mice with or without CTX injury **(Fig. 6)**. Consistent with the *in vitro* and *in vivo* observations in Fig. 4 and 5, an increase in AT2R protein expression was not detected in SCs harvested from noninjured muscles **(Fig. 6A, B)**, whereas AT2R mRNA levels were increased >70 fold in these cells **(Fig. 6C)**. Also, consistent with the results in Fig. 4 and 5, AT2R protein levels were increased in SC-CAG-AT2R-derived SCs 5 days after CTX injury **(Fig. 6A, B)**. This increase in AT2R protein was associated with increased p-GSK3β^Ser9^, p-Akt^Ser473^, myogenin and desmin expression. qRT-PCR from the same set of cells showed increased expression of Myog, Mymk, Mymx, follistatin and Myh3, whereas Pax7 and MyoD expression was not altered **(Fig. 6C)**. TOPFlash reporter assay showed an increase in TCF/LEF promoter activity during CTX-induced regeneration, and this effect was potentiated by AT2R overexpression, indicating an activation of the GSK3β/β-catenin pathway in SCs of SC-CAG-AT2R mice **(Fig. 6D)**.

**Fig. 6.**
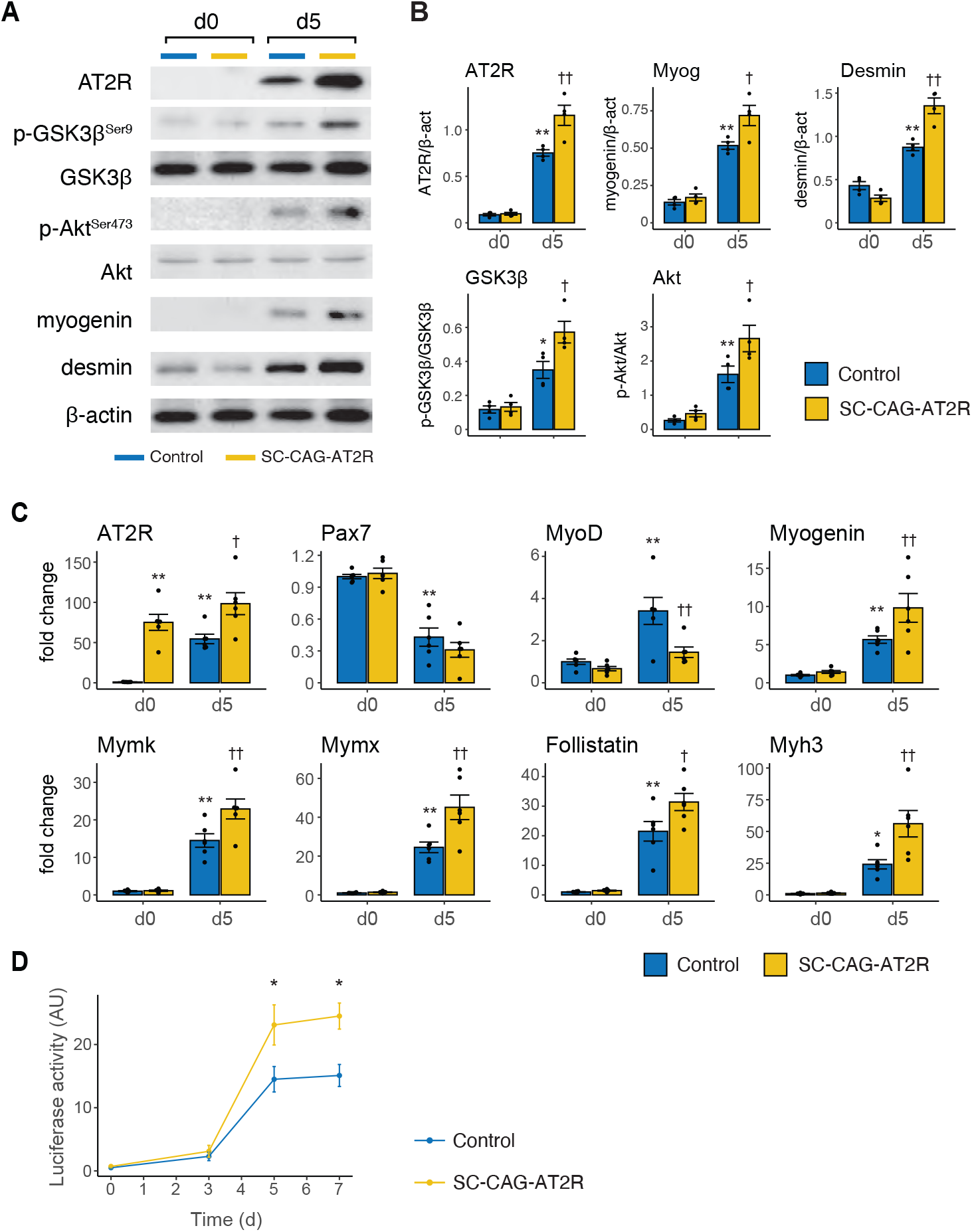
AT2R overexpression in SCs potentiates SC differentiation *in vivo*. **(A, B)** CTX was injected into TA muscles of tamoxifen-administered SC-CAG-AT2R and control (SC-EYFP) mice, and SCs were collected from these mice before (d0) and 5 days after injury (d5). Expression of AT2R, p-GSK3β^Ser9^/GSK3β, p-Akt^Ser473^/Akt, myogenin and desmin was analyzed by immunoblotting. Immunoblot results were quantified in (B). **(C)** From the same set of samples as in (A), gene expression was analyzed by qRT-PCR. In (B) and (C), asterisks (*) and daggers (†) indicate statistical significance compared to d0-control and d2-control group, respectively. N=4 in (B) and N=6 in (C); mean±SEM; *p<0.05; **p<0.01; †p<0.05; ††p<0.01. **(D)** TopFlash reporter vector was electroporated into TA muscles of SC-CAG-AT2R and control SC-EYFP mice, and CTX was injected 2 weeks after electroporation. Muscles were harvested at the indicated time points, and luciferase activity was measured. Asterisks (*) indicate statistical significance between SC-CAG-AT2R and control mice for each time point. N=4; mean±SEM; *p<0.05.

### Protection of SC-CAG-AT2R mice from Ang II-mediated inhibition of skeletal muscle regeneration

We have demonstrated that high Ang II levels following Ang II infusion in wildtype mice and possibly in MI-induced CHF animal models inhibit skeletal muscle regeneration, and this inhibition is associated with blunted AT2R induction in SCs. To determine whether an increase in AT2R expression prevents Ang II-mediated inhibition of skeletal muscle regeneration, we infused SC-CAG-AT2R or control mice with Ang II after CTX injury **(Fig. 7)**. qRT-PCR of whole muscle samples showed an additive increase in AT2R at d5 in sham-infused mice **(Fig. 7A)**. However, Ang II suppressed endogenous AT2R mRNA expression, and this inhibition was partially restored in SC-CAG-AT2R mice. Expression of Myog, Mymk, Mymx and Myh3 was also partially restored in SC-CAG-AT2R mice infused with Ang II, suggesting that Ang II-mediated inhibition of muscle regeneration could be prevented by AT2R overexpression. Consistently, myofiber cross sectional area of regenerating myofibers was restored in SC-CAG-AT2R mice infused with Ang II **(Fig. 7B)**. Furthermore, TOPFlash reporter assay showed that TCF/LEF reporter activity was blunted by Ang II infusion, but was restored in SC-CAG-AT2R mice **(Fig. 7C)**.

**Fig. 7.**
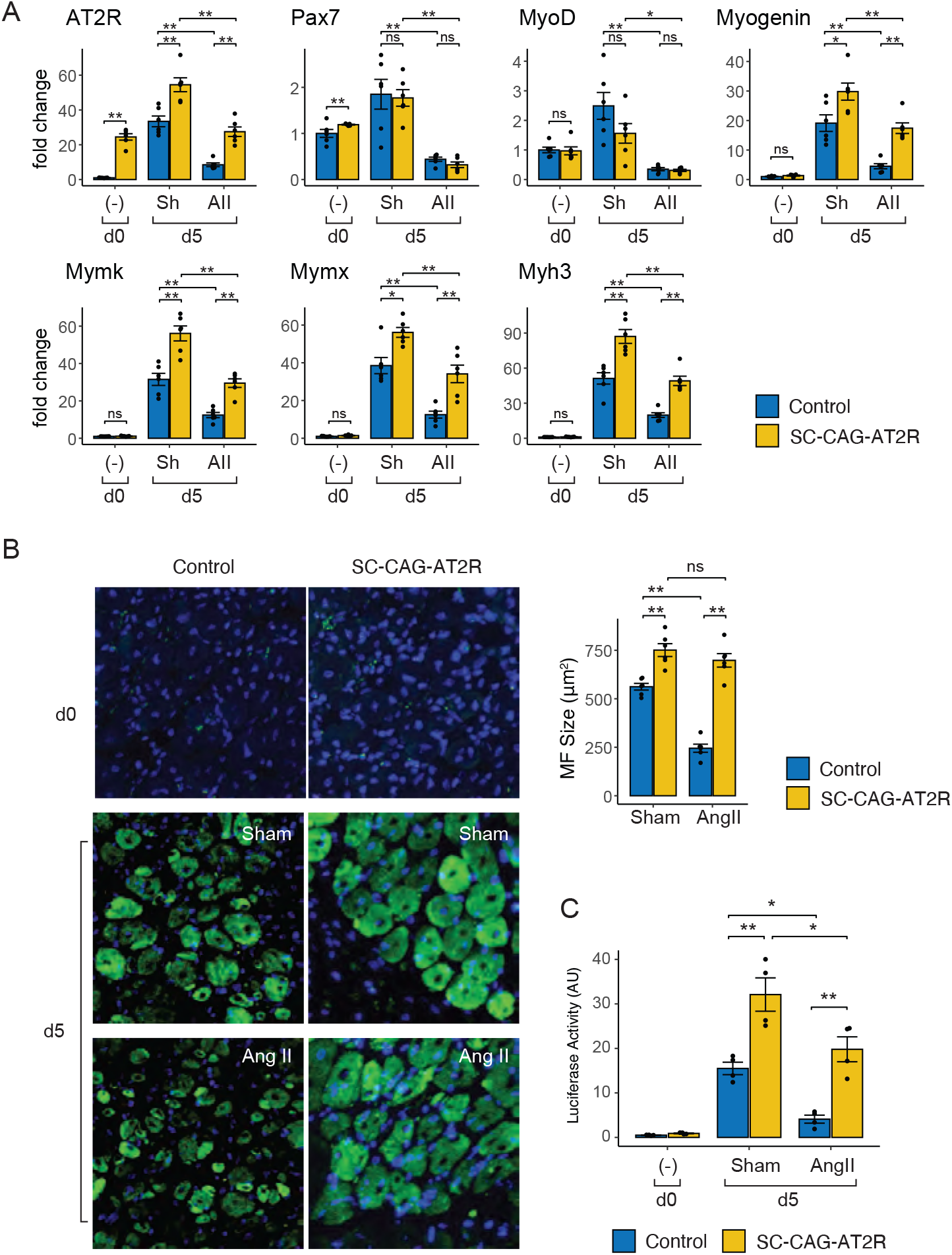
AT2R overexpression in SCs protects against Ang II-mediated inhibition of SC differentiation *in vivo*. **(A)** After tamoxifen administration, SC-CAG-AT2R and control SC-EYFP mice were injected with CTX into hindlimb muscles, and infused with Ang II via osmotic minipumps. Muscles were harvested at d5, and gene expression before (d0) and after (d5) injury in TA muscles was quantified by qRT-PCR. N=6; mean±SEM; *p<0.05; **p<0.01. **(B)** From the same set of samples as in (A), newly generated myofibers were visualized by EYFP fluorescence. Cross sectional area (CSA) of EYFP+ myofibers was quantified (right panel). N=6; mean±SEM; *p<0.05; **p<0.01. **(C)** TopFlash reporter vector was electroporated into TA muscles of SC-CAG-AT2R and control SC-EYFP mice after tamoxifen administration. 2 weeks after electroporation, CTX injection and Ang II infusion were performed as in (A), and muscles were harvested before (d0) and after (d5) the injury. Luciferase activity was measured and normalized by *Renilla* luciferase activity from pGL4.74[hRluc/TK] vector. N=4; mean±SEM; *p<0.05; **p<0.01.

## Discussion

Chronic diseases like advanced CHF and CKD are characterized by increased systemic Ang II levels and cachexia. We previously demonstrated that Ang II infusion in rodents results in skeletal muscle wasting and reduced skeletal muscle regeneration. Muscle SCs are the muscle stem cells that contribute to muscle growth and repair after injury and during development. Ang II signals mainly via the AT1R and AT2R, and we have previously shown that AT2R expression is markedly increased during SC differentiation. In this study, we found that the GSK3β/β-catenin pathway plays an important role downstream of AT2R, leading to activation of myogenic genes. Using the phosphoprotein array, we found that GSK3β/β-catenin was one of the top canonical pathways that were altered in SCs following AT2R knockdown **(Fig. 1)**. The GSK3β/β-catenin pathway is activated (increased GSK3β^Ser9^ phosphorylation and TCF/LEF transcriptional activity) during SC differentiation, and AT2R silencing blunted this increase, leading to inhibition of SC differentiation **(Fig. 2A-D)**. Interestingly, while GSK3β inhibitor restored SC GSK3β/β-catenin activity and SC myogenic capacity in SCs with AT2R knockdown, Wnt3A did not, suggesting that AT2R regulates GSK3β/β-catenin pathway independently and/or in parallel with the canonical Wnt signaling **(Fig. 3)**. To further investigate the role of AT2R in SC differentiation, we generated transgenic mice that overexpress AT2R specifically in SCs (SC-CAG-AT2R mice), and found that AT2R overexpression enhances their differentiation **(Fig. 4)**. Interestingly, however, we found that AT2R protein levels were not altered in undifferentiated SCs in SC-CAG-AT2R mice likely due to post-transcriptional suppression **(Fig. 4C, F)**. Indeed, AT2R overexpression did not induce SC differentiation *in vitro* or activate skeletal muscle regeneration *in vivo* without an activation cue (e.g. switching to differentiation medium or following muscle injury), suggesting that AT2R protein expression is strictly regulated post-transcriptionally to prevent premature initiation of SC differentiation processes. We have previously shown that systemic Ang II infusion prevented skeletal muscle regeneration *in vivo*, which was associated with inhibition of AT2R increase. We found that SC-CAG-AT2R mice were at least partially protected against Ang II-mediated reduction of skeletal muscle regeneration **(Fig. 8)**, further supporting the role of AT2R in the regulation of SC differentiation and skeletal muscle regeneration. Together, these results suggest that the AT2R/GSK3β/β-catenin signaling pathway could serve as a potential therapeutic target to counteract lower muscle regenerative capacity during chronic disease conditions, like CHF and CKD, that are characterized by heightened activation of the RAS.

The Wnt signaling pathway plays a key role in many aspects of development, tissue homeostasis, stem cell fate, organogenesis, cancer and tissue fibrogenesis. Wnt ligands bind to membrane receptor co-complexes with Frizzled (Fzd) and low-density lipoprotein receptor protein (LRP) families. In the absence of a Wnt stimulus, β-catenin is a target for UPS-mediated degradation^71,72^. Canonical Wnt signaling suppresses β-catenin ubiquitylation and degradation, and the accumulated β-catenin translocates to the nucleus, where it binds to TCF/LEF transcription factors and activates expression of multiple TCF/LEF target genes^73–75^. Canonical Wnt ligands are shown to promote myoblast differentiation *in vitro* and can increase the expression or activity of MyoD and/or myogenin. Moreover, β-catenin can interact with MyoD and enhance its ability to bind and activate target genes. However, conflicting results were reported to describe the role of TCF/LEF. For example, Kim *et al*.^76^ showed that β-catenin knockdown impairs myoblast differentiation, whereas dominant-negative forms of TCF/LEF did not, suggesting that β-catenin functions independent of TCF/LEF during myogenesis. In contrast, Agley *et al*.^77^ showed that dominant-negative TCF4 in human adult primary myoblasts impaired differentiation. Nonetheless, these studies support the notion that β-catenin signaling plays an important role in myoblast differentiation. It is possible that AT2R regulates TCF/LEF-independent functions of β-catenin in addition to TCF/LEF transcriptional activation **(Fig. 2C, 3E, 6D)**.

It is important to note that Wnt signaling seems to act as a double-edged sword: on the one hand it induces myogenic differentiation of SCs during regeneration, or it can repress the myogenic lineage and forces SCs to an alternative lineage, leading to fibrosis in an aged muscle^68,78^. Myoblast differentiation is correlated with upregulation of canonical Wnt signaling, and ectopic Wnt induces premature muscle differentiation whereas Wnt inhibition interferes with muscle differentiation^68^.

These reports suggest that Wnt signaling is strictly regulated in SCs to prevent ectopic/premature myogenesis. Our data show that AT2R activates the GSK3β/β-catenin pathway downstream of, or independent of Wnt: GSK3β inhibitor BIO or forced expression of caAkt restored myogenic capacity in AT2R silenced SCs, whereas Wnt3A did not **(Fig. 3)**. In Barx2 deleted SCs **(Fig. 2E-G)**, the expression of canonical Wnt ligands (Wnt7b and Wnt10b) were increased together with Pax7 and MyoD, whereas non-canonical Wnt ligand (Wnt5a) expression was decreased. The association of AT2R expression with the canonical Wnt ligands, Pax7 and MyoD suggests that AT2R acts upstream of the Barx2/β-catenin complex **(Fig. 2G)**.

Given the variety of Wnt ligands and the diversity of Wnt signaling pathways^79^, it is possible that the synthesis of different Wnt ligands and activation of their downstream signaling intermediates are precisely regulated during SC differentiation processes to prevent unwanted outcomes (e.g. premature differentiation or fibrosis development). Interestingly, we found that transgenic overexpression of AT2R in SCs (SC-CAG-AT2R mice) did not increase AT2R protein levels, whereas its mRNA expression was markedly increased **(Fig. 4C, F)**. Similarly, AT2R protein expression was not increased following transfection with the AT2R plasmid vector in cultured SCs, C2C12 myoblasts, or other non-myogenic cell lines such as HeLa, HEK293 or CHO, even though AT2R mRNA expression was markedly increased (data not shown). In contrast, AT2R protein was increased in SC-CAG-AT2R-derived SCs after cells were induced to differentiate **(Fig. 4F and 6A)**, suggesting that AT2R protein expression is post-transcriptionally regulated in SCs. Detectable levels of AT2R protein were observed in SCs only after cells were induced to differentiate **(Fig. 4F and 6A)**, suggesting an unknown post-transcriptional mechanism whereby AT2R protein levels are suppressed in undifferentiated SCs. Importantly, AT2R mRNA was expressed at very low levels in undifferentiated SCs, but markedly upregulated upon induction of SC differentiation. These data suggest that AT2R expression is strictly regulated via both transcriptional and post-transcriptional mechanisms. It is possible that this strict AT2R regulation plays a role in controlling Wnt/GSK3β signaling in SCs, likely preventing premature SC differentiation.

Backlund *et al*. have shown post-transcriptional regulation of AT1R via binding of GAPDH protein to the 3’-UTR of AT1R mRNA^80^. Also, computational models such as TargetScan^81^ or miRDB^82^ predict multiple putative miRNA target sites in AT2R 3’-UTR (data not shown). However, it is unlikely that these mechanisms are responsible for post-transcriptional regulation of AT2R in SCs, as the AT2R transgene in SC-CAG-AT2R mice does not contain 3’-UTR sequence of the AT2R gene **(Fig. S2A)**. It is also possible that AT2R protein levels are affected by proteins that directly bind and regulate its expression (e.g. via regulation of protein stability). In addition to the well-established AT1R and AT2R protein interaction, several proteins are shown to directly bind to AT2R, including microtuble associated tumor suppressor 1 (MTUS1)^83^, epidermal growth factor receptor 3 (ErbB3)^84^, Src homology region 2 domaincontaining phosphatase-1 (SHP-1)^85^, and promyelocytic leukemia zinc finger (PLZF)^86^, although none of these proteins are known to function in SCs nor to regulate AT2R protein expression. Further studies are required to elucidate the post-transcriptional regulation of AT2R expression.

Cell fate choice between self-renewal and differentiation is critical for SCs to regulate appropriate muscle regenerative processes. The onset of differentiation is due to a transition from Notch signaling to Wnt signaling in SCs, and their crosstalk is mediated by GSK3β; Notch maintains GSK3β in an active form and this effect is inhibited by canonical Wnt ligands^68^. We have previously shown that AT1R expression in SCs is decreased at the onset of differentiation, and AT1R signaling inhibits Notch activation^41^. These reports together with results from our current study suggest that both AT1R and AT2R coordinately regulate SC proliferation and differentiation processes via GSK3β. It is important to note that circulating Ang II levels are increased^87–89^ and AT2R induction in SCs is transcriptionally suppressed^43^ in CHF conditions, possibly leading to inhibition of both Notch as well as Wnt pathways. A dysregulation in these pathways is likely an underlying cause of decreased SC function and muscle regenerative capacity in these animals.

The AT2R regulates various cellular processes via different downstream signaling pathways in a context dependent manner. Functions of AT2R include vasodilation through nitric oxide (NO) and cGMP stimulation, natriuresis, angiangiotenesis, antiproliferation, and decreased fibrosis. These effects are observed in various cell types and tissues, including endothelium, vascular smooth muscle, heart, brain and kidney^90^. AT2R is also known to interact with other receptors, such as bradykinin receptor, NGF receptor and PDGFR^91^. NO has been shown to mediate SC activation, including morphological hyptertrophy and decreased adhesion in the fiber-lamina complex^92^. NO release is triggered by mechanical stretch of skeletal muscle, leading to activation of SCs, and NO treatment has been suggested to enhance the effectiveness of exercise, especially in the elderly^93^. Although it remains to be determined whether AT2R regulates NO production in SCs, it is interesting to note that GSK3β has been shown to increase the expression of inducible nitric oxide synthase (iNOS) and NO production in macrophages^94^, and to regulate IL-6-mediated increase in iNOS and NO in microglia^95^.

Interestingly, our data showed that AT2R overexpression increases SC differentiation *in vitro* without supplementation of its ligands in culture **(Fig. 4C-F)**. Although we have shown that SCs express various RAS components including angiotensinogen, renin, ACE and ACE2, it is not known whether angiotensin peptides are locally produced during SC differentiation and muscle regeneration. Since AT2R binds to other angiotensin peptides, including angiotensin III (Ang III)^96^ and Ang(1-9)^97^, these peptides might be involved in AT2R signaling in SCs. It is also plausible that AT2R elicited the observed effects independent of ligand binding: In contrast to the conventional ligand-receptor regulation, stimulation of AT2R by Ang II is not required for induction of apoptosis but dependent on the level of AT2R protein expression^98^. From a clinical perspective, the absence of a requirement for additional ligand supplementation in order to stimulate AT2R signaling *in vivo* is critical to avoid potential side effects associated with ligands.

In summary, our data demonstrate that the AT2R is a novel upstream signaling pathway that regulates GSK3β/β-catenin activity in SCs, and plays a critical role in SC differentiation and potentially fibrosis development. Activation of AT2R signaling by increasing its protein expression could be a novel therapeutic strategy to potentiate SC differentiation capacity in patients with lowered muscle regeneration such as patients with CHF, CKD or aging.

---

**Fig. S1.**
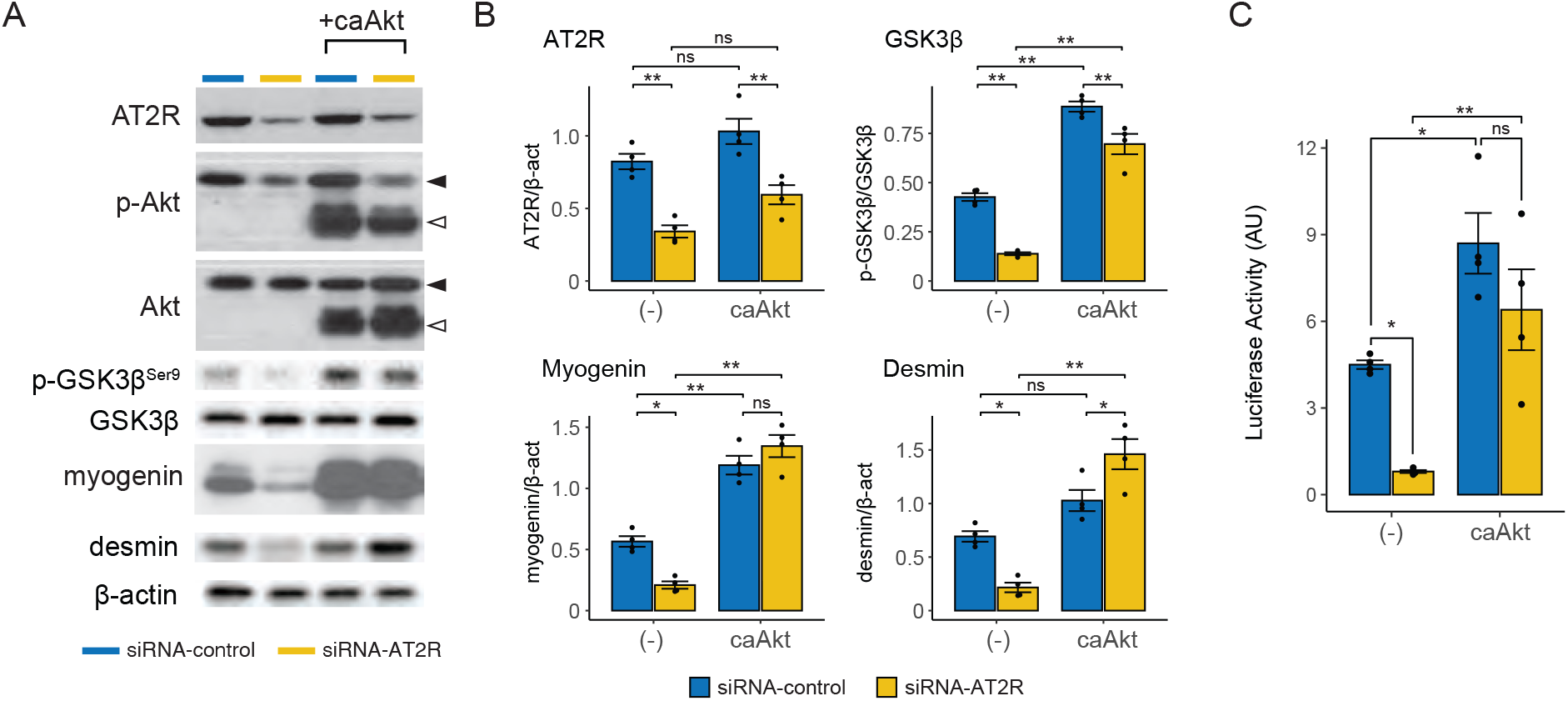
Constitutively-active Akt inhibits GSK3β pathway in AT2R knocked down SCs. **(A)** Immunoblot analysis of Akt/GSK3β pathway and myogenic differentiation markers. Primary SCs were transfected with AT2R or control siRNA, induced to differentiate for 2 days, and protein expressions were analyzed by immunoblotting. **(B)** Quantification of the immunoblots from (A). N=4, mean±SE, *p<0.05, **p<0.01. **(C)** Quantification of TCF/LEF reporter activity. Primary cultured SCs were transfected with TOPFlash reporter vector M50 or mutant control vector M51, together with AT2R or control siRNA. Cells were induced to differentiate, and luciferase activity was measured at the indicated time points. Firefly luciferase activity from M50/M51 vectors was normalized by *Renilla* luciferase activity from the control pGL4.74[hRluc/TK] vector. N=4; mean±SEM; *p<0.05; **p<0.01.

**Fig. S2.**
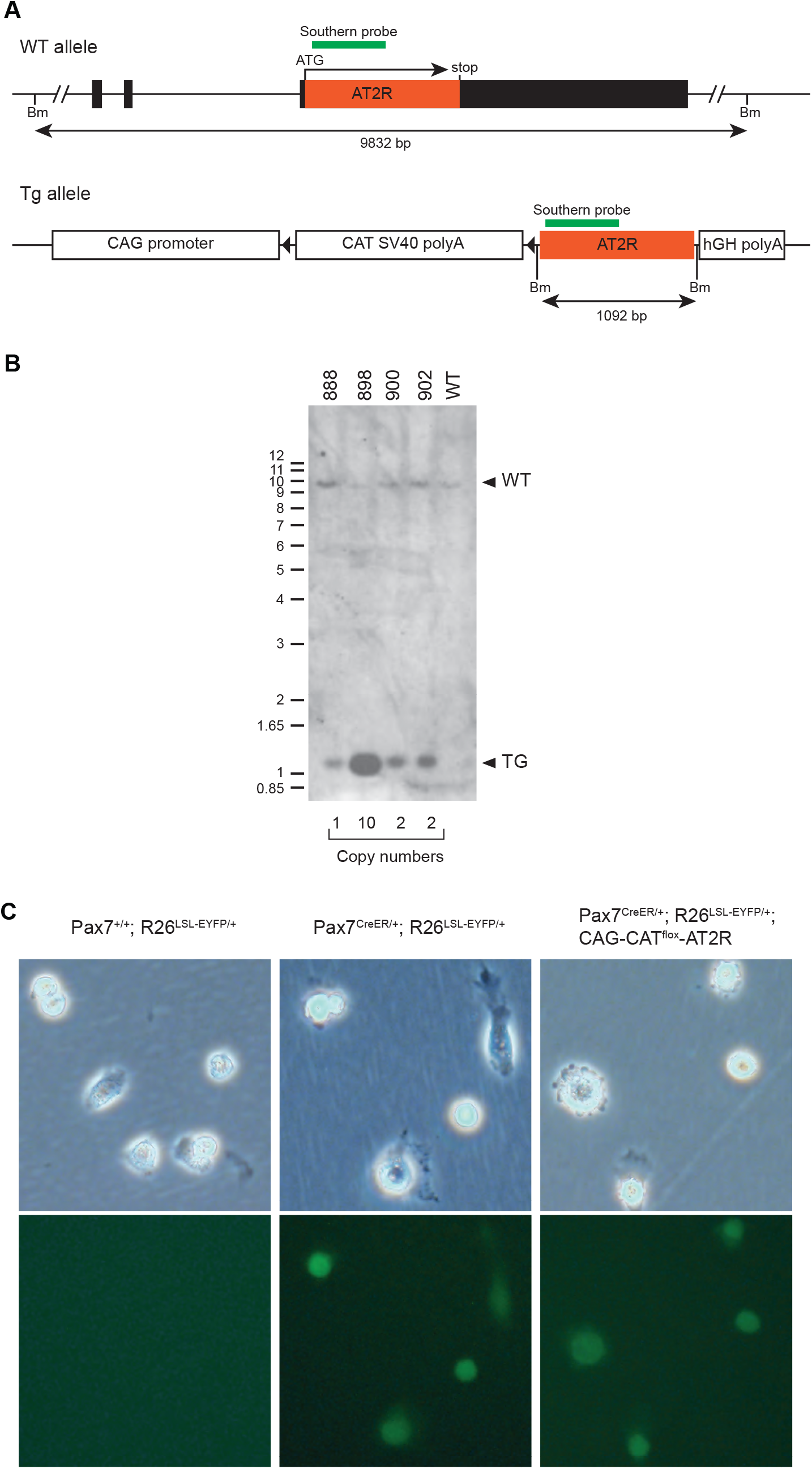
Generation of SC-CAG-AT2R mice. **(A)** Map of mouse AT2R gene (top) and transgenic vector for CAG-CAT^flox^-AT2R mice (bottom). **(B)** Southern blotting of AT2R gene in four CAG-CAT^flox^-AT2R founder lines (lines 888, 898, 900 and 902) with the probe indicated in (A). Genomic DNA of these animals were digested with BamHI. The location of the restriction enzyme digestion sites and the predicted DNA fragment sizes are shown (A). Copy numbers from each line are shown at the bottom. **(C)** Primary SCs were collected from SC-CAG-AT2R, SC-EYFP and negative control Pax7^+/+^; R26^LSL-EYFP/+^ mice, and bright field and EYFP fluorescence images are shown.

**Fig. S3.**
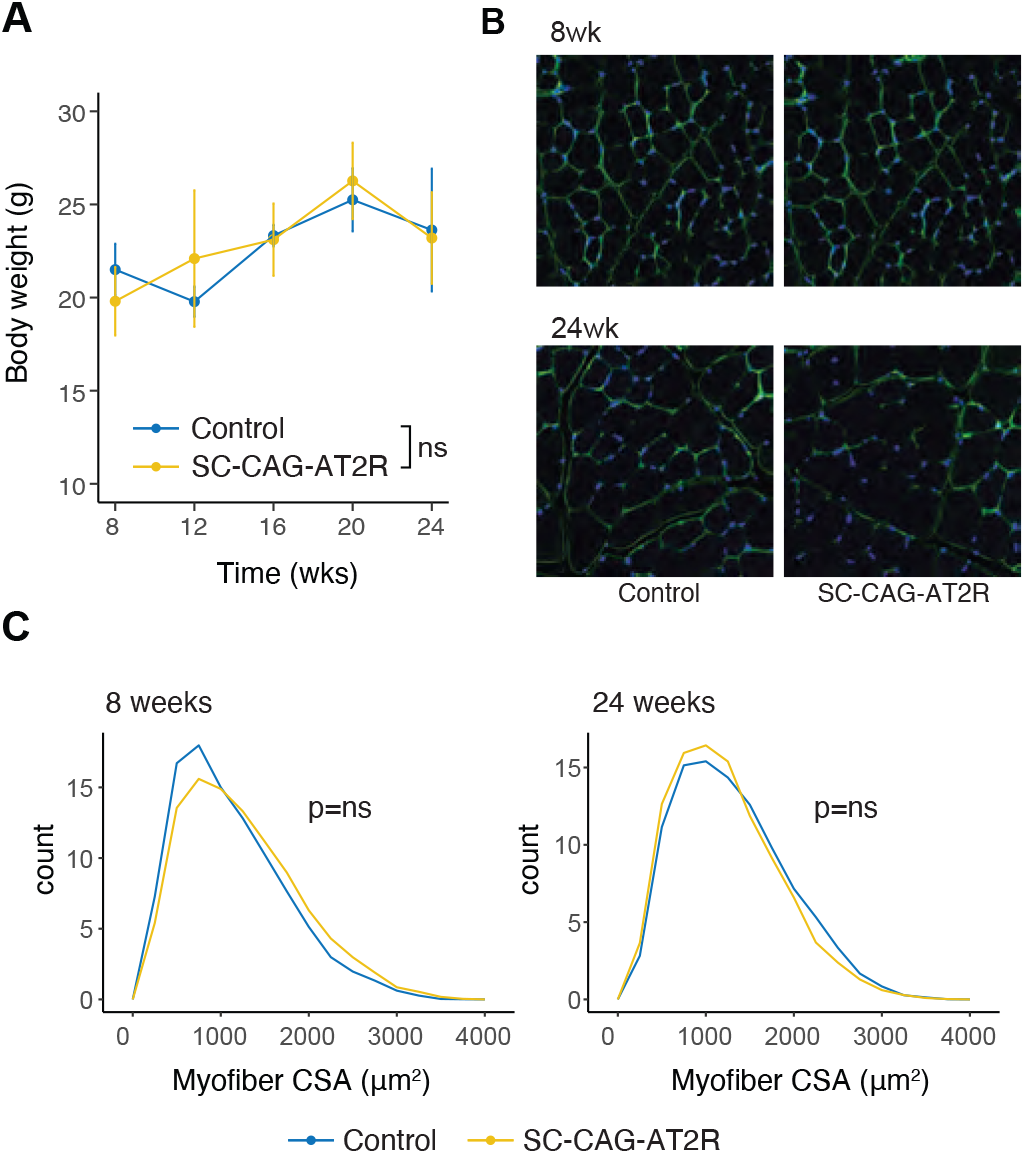
Body weight and myofiber size in SC-CAG-AT2R mice. **(A)** Tamoxifen was administered to SC-CAG-AT2R and control (Pax7CreER/+; R26^LSL-EYFP^) mice at 6 weeks of age, and body weight changes were measured during 8 to 24 weeks. N=6; mean±SEM. **(B)** From the same set of animals as in (A), tibialis anterior muscle sections were stained for laminin (B), and myofiber cross sectional area was measured (C).

**Sup Table 1 Phosphoarray raw data**

Calculated intensity values of PhosphoExplorer Antibody Array. For each spot on the array, median signal intensity is extracted from array image, and the average signal intensity of the replicate spots were determined for each antibody. Within each array slide, the mean value of the average signal intensity for all antibodies on the array was determined, and the signal was normalized by the median signal. Normalized data were further calculated to determine the log fold change values between AT2R siRNA-treated and control siRNA-treated samples.

## Data Availability Statement

The data, materials and animals that support the findings of this study are available from the corresponding author upon reasonable request.

## Author Contributions

C.S.L: Collection and assembly of data, data analysis and interpretation, and manuscript writing. B.C: Conception and design, and interpretation of data. P.D. Conception and design, interpretation of data, financial support, and manuscript writing. T.Y: Collection and assembly of data, data analysis and interpretation, financial support, and manuscript writing.

## Acknowledgement

The study was supported by grants from American Heart Association (19TPA34850165 and 15SDG25240022 for TY), US Department of Veterans Affairs (VA-I01-BX004220 and VA-IK6BX004016 for BC) and NIH/NHLBI (R01HL070241 and R01HL070241-AS for PD). We thank Daniel Davis (University of Missouri Animal Modeling Core) for assistance with generation of transgenic animals, Dina Guapp (Tulane Histology Core) for assistance with tissue sectioning and Alan Tucker (Tulane Flow Cytometry and Cell Sorting Core) for assistance with cell sorting.

